# Micro-aeration-enhanced Anaerobic Digestion for the Stabilization of Coffee-Processing Wastewater

**DOI:** 10.1101/2025.06.09.658626

**Authors:** Kayode J. Taiwo, Samuel O. Ogundipe, William L. Kerr, Ronald B. Pegg, Joon Hyuk Suh, Joseph G. Usack

## Abstract

Coffee processing wastewater (CPW), a byproduct of agro-industrial operations, contains high organic loads alongside recalcitrant and potentially inhibitory compounds such as caffeine and tannins. This study evaluated the performance of micro-aeration-enhanced anaerobic digestion (MA-AD) for the treatment and valorization of CPW to promote a more sustainable approach to coffee production. Oxygen was intermittently introduced via oxidation-reduction potential-controlled dosing, allowing for comparative assessment across anaerobic and micro-aerobic redox regimes. While both conventional anaerobic digestion (AD) and MA-AD achieved comparable reductions in total and volatile solids (>48% and >60%, respectively) and total and soluble chemical oxygen demand (>66% and >86%, respectively), MA-AD exhibited significantly higher total suspended solids concentrations and turbidity in later phases, likely due to gas sparging-induced floc disruption and particulate release. pH profiles indicated a shift toward increased acidification under MA-AD, without compromising process stability, with both reactors stabilizing between pH 6.8–7.1. Caffeine degradation was accelerated under MA-AD in the first dosing phase (>85% removal in 28 h), though long-term degradation efficiency converged with the control. Methane production was consistently lower in MA-AD (up to 43% reduction), attributed to the oxygen sensitivity of methanogens and possible substrate competition. These results underscore the importance of oxygen dose regulation, redox control, and microbial adaptation in optimizing MA-AD performance. The findings support MA-AD as a promising strategy for enhancing hydrolysis and partial removal of recalcitrant compounds in CPW. However, further refinement is required to sustain biogas quality and yield at scale.

## 1.0 Introduction

Coffee is one of the most widely traded global commodities, with trade volumes expected to exceed 10.5 million tons in 2024/2025 (USDA, 2024). Globally, it ranks as the second most valuable commercial product after oil, with significance across diverse economies and sectors (Ponte, 2002). The coffee processing industry plays a substantial role in driving economic growth at both national and international levels (Rattan et al., 2015). Also, about one-third of the global population consumes coffee, reflecting its pervasive socio-cultural impact. As the global coffee market expands, consumer expectations for fair and sustainable sourcing have also increased. However, coffee production practices frequently do not align with these expectations in many countries, primarily due to limited regulatory oversight, inadequate incentives, gaps in knowledge, and a lack of resources needed to support sustainable practices (Dadi et al., 2018). A direct consequence of such non-sustainable practices is the discharge of untreated coffee processing wastewater (CPW) into surrounding freshwater bodies, adversely affecting aquatic ecosystems and the health of those whose livelihoods depend on them. Meanwhile, in high-consumption areas such as the United States, the United Kingdom, and Europe, coffee brewing practices are resource-intensive, requiring 42-57 grams of roasted ground coffee per liter of water, respectively (IARC, 1991; ICO, 2011), underscoring the environmental burden across the coffee supply chain.

Similarly, CPW, which is a byproduct of coffee bean production and processing, is characterized by a high organic load (i.e., high biochemical oxygen demand (BOD) and chemical oxygen demand (COD) (Chen et al., 2018). CPW contains a large amount of organic matter, including proteins, sugars, caffeine, tannins, and other polyphenolic compounds. These compounds, particularly tannins, polysaccharides, and their mixtures, contribute to the dark coloration of the effluent due to the presence of melanoidins, which are resistant to biological degradation (Cárdenas et al., 2009; Ijanu et al., 2020; Zayas et al., 2007). When released into aquatic environments, these substances promote the formation of anaerobic conditions, fostering the proliferation of microbial organisms. This microbial activity depletes dissolved oxygen (DO) levels, disrupts aquatic food chains, and generates toxic byproducts, posing severe ecological risks to aquatic ecosystems (Singh et al., 2021). Furthermore, contamination of drinking water sources by these microorganisms can lead to significant health risks for humans, such as skin irritation and gastrointestinal disturbances (Figueroa Campos, 2022).

In addition, CPW contains a high concentration of suspended solids, which increases turbidity in water bodies (Ijanu et al., 2020; Novita et al., 2012). This turbidity contributes to opacity, blocking light penetration and reducing photosynthesis in aquatic plants, further disrupting the balance of aquatic ecosystems (Ijanu et al., 2020; Zayas et al., 2007; Takashina et al., 2018; Tokumura et al., 2006). These factors collectively make CPW a significant environmental pollutant, requiring advanced treatment methods to mitigate its impact on aquatic ecosystems. As global coffee consumption increases, addressing coffee wastewater management has thus become essential to comply with rising environmental standards and minimize the industry’s ecological footprint.

Traditional treatment methods often struggle with CPW’s complex and highly variable composition, necessitating innovative approaches such as anaerobic digestion (AD) and micro-aeration-assisted anaerobic digestion (MA-AD) (Ijanu et al., 2020). AD is a widely adopted biological process for treating high-strength organic wastewater, including agricultural, food processing, and industrial effluents (del Agua et al., 2015; Zhou et al., 2024). AD involves the breakdown of organic matter by microorganisms in the absence of oxygen, producing biogas composed primarily of methane (50-65%) and carbon dioxide (35-50%) (Angenent et al., 2022; Zhou et al., 2024). The conversion of organic substrates into biogas proceeds through four trophic stages: hydrolysis, acidogenesis, acetogenesis, and, finally, methanogenesis (Laiq Ur Rehman et al., 2019). The AD process is known for its ability to reduce the organic load of wastewater while simultaneously generating renewable energy in the form of biogas. However, the efficiency of AD is influenced by various factors, which can either improve or inhibit the process. These factors include the composition of the substrate, organic loading rate (OLR), mixing conditions, presence of inhibitory compounds, temperature, pH levels, carbon-to-nitrogen ratio, microbial diversity, and retention time (Gyadi et al., 2024).

The high levels of inhibitory compounds in CPW, such as phenols, caffeine, and tannins, can impede microbial activity, leading to lower biogas yields and incomplete degradation of organic matter (Figueroa Campos, 2022). In addition, CPW’s low alkalinity, acidic pH, and lack of essential nutrients and trace elements further complicate its treatment through AD (Ijanu et al., 2020). Several studies have reported low biogas yields, COD removal, and the accumulation of volatile fatty acids (VFAs) during mono-digestion of CPW. Qiao et al. (2013) further reported reactor failures under such conditions. However, Du et al. (2020) noted that process failure could be mitigated and biogas production enhanced by supplementing trace minerals. Chen et al. (2018) and Selvamurugan et al. (2010) suggested using co-digestion as a strategy to stabilize the AD treatment of coffee residue. Still, the effectiveness in breaking down biologically recalcitrant compounds remains almost the same as the mono-digestion of CPW. Even with high-rate anaerobic reactors such as upflow anaerobic sludge blanket reactors, AD alone is often insufficient for achieving complete organic matter removal. A considerable portion of recalcitrant compounds, such as lignin, tannins, phenols, and humic acids, remain in the treated effluent, as they are highly resistant to biological degradation (Bruno & Oliveira, 2013; Campos et al., 2013; Gomes de Barros et al., 2020; Zayas et al., 2007; Villa-Montoya et al., 2017).

Furthermore, anaerobic microbiomes show inherent limitations in degrading recalcitrant compounds, particularly those with aromatic ring structures, due to the absence of oxygen as a strong electron acceptor and the lack of robust enzymes. Aerobic systems, by contrast, utilize oxygen and powerful enzymes to cleave such molecular bonds, making them significantly more effective in breaking down these complex compounds (Ortiz-Ardila et al., 2024). Non-biological oxidative methods have, however, proven effective. For example, Tokumura et al. (2006) reported nearly complete decolorization of coffee processing effluent using the photo-Fenton process (UV/Fe² /H O). Similarly, Takashina et al. (2018) demonstrated the effectiveness of a combined ozone and UV treatment (O /UV), achieving 98% removal of caffeine and 99% degradation of colorizing compounds in synthetic coffee wastewater. Furthermore, Yamal-Turbay et al. (2012) reported 100% removal of caffeine using the photo-Fenton method combined with hydrogen peroxide. However, while effective, these methods are often associated with high operational costs and technical complexity due to the need for specialized equipment, energy-intensive processes, and expensive chemicals, making them less sustainable (Gogate & Pandit, 2004). Therefore, to overcome the challenges associated with conventional anaerobic systems, aerobic systems, and non-biological oxidative methods, relatively low-cost modifications, such as micro-aeration, need to be explored.

Micro-aeration offers a cost-effective and sustainable alternative, leveraging natural microbial processes with minimal infrastructure modifications and lower energy requirements, making it more practical and accessible, particularly in resource-limited settings (Rajagopal & Goyette, 2024). Unlike traditional AD processes, which operate under strict anaerobic conditions, micro-aeration allows for the controlled dosing of oxygen at low levels to promote the growth of facultative microorganisms that augment the breakdown of particulate and recalcitrant compounds. Studies have also shown that micro-aeration can enhance biogas yield, COD removal, promote microbial diversity, and reduce sludge production in the treatment of various organic waste streams (Ding et al., 2024; Fu et al., 2023; Xu et al., 2021). Additionally, it has been shown to prevent acidification, enhance hydrolysis, mitigate process instability, reduce hydrogen sulfide accumulation, and stimulate microbial interactions, further optimizing AD performance (Barati Rashvanlou et al., 2020; Li et al., 2024; Morais et al., 2024; Song et al., 2020; Zhan et al., 2023; Zhu et al., 2022). Furthermore, Fu et al. (2023), Magdalena et al. (2022), Nguyen and Khanal (2018), and Shrestha et al. (2017) reported enhanced degradation of various recalcitrant compounds using micro-aeration in other AD applications. Still, to our knowledge, no research has evaluated its effectiveness in treating coffee processing residues.

The synergy between micro-aeration and AD offers a promising solution for treating CPW. This hybrid approach can address the limitations of conventional anaerobic reactors, such as slow degradation rates and sensitivity to recalcitrant compounds, by creating a more robust microbial community. Integrating micro-aeration in full-scale anaerobic reactors poses technical challenges, including maintaining stable conditions and achieving uniform oxygen distribution, which is critical for consistent treatment performance (Botheju & Bakke, 2011). Factors such as the dosage and timing of oxygen injection, reactor configuration, and microbial community dynamics play a crucial role in determining the efficiency of the process (Botheju & Bakke, 2011; Chen et al., 2020). Achieving the right balance between anaerobic and aerobic conditions is essential, as excessive oxygen can inhibit methanogenesis and shift the microbial community towards aerobic degradation pathways. At the same time, too little oxygen may have no significant effect. Thus, understanding the interactions between oxygen levels, microbial activity, and degradation of complex organic matter is critical to successfully implementing this technology at scale.

This study aims to address these knowledge gaps by evaluating the performance of MA-AD in treating CPW. Specifically, the study focuses on optimizing oxygen dosage and assessing the impact of micro-aeration on methane yield, COD, solids, and turbidity reduction, as well as the degradation of recalcitrant compounds. By providing insights into the underlying mechanisms and operational conditions, this research seeks to contribute to the broader goal of sustainable wastewater management in the coffee industry.

## 2.0 Materials and methods

### 2.1 Reactor set-up

The reactors consisted of two benchtop fermenters (BioFlow3100, New Brunswick Scientific Co., Edison, NJ, USA), each with a 1.5-L vessel and an operational volume of 1 L. Each reactor was outfitted with a headplate that included a sampling port, a gas sparger, an oxygen-reduction potential (ORP) probe (Sensorex, Garden Grove, CA, USA), an influent inlet, and an effluent outlet. The reactors were set up as described by Usack et al. (2012) and operated as continuous stirred tank reactors (CSTR) using an overhead stirrer (Fisherbrand Compact Digital, Waltham, MA, USA) integrated with an ORP controller (Oakton 220, Vernon Hills, IL, USA). A 3-L water bath (Fisher Scientific, Hampton, NH, USA) maintained the reactor temperature at 37 ±1°C. The effluent was withdrawn using a peristaltic pump (Masterflex®, Vernon Hills, IL, USA) for further analysis. Biogas production was monitored with integrated flow meters (BPC Instruments, Mobilvägen, LU, Sweden). The reactors comprised 1) a control reactor system for conventional AD and 2) an experimental reactor system for MA-AD **(Figure 1A and 1B)**. The reactors were allowed to reach pseudo-steady-state operation before initiating the experimental phase of the study. During the experimental phase, one reactor was sparged with oxygen to induce microaerobic conditions, while the other was supplied with 100% nitrogen gas (N) to maintain AD conditions. The ORP in the micro-aeration reactor was regulated using integrated ORP controllers following the guidelines of Nguyen and Khanal (2018).

**Figure 1:**
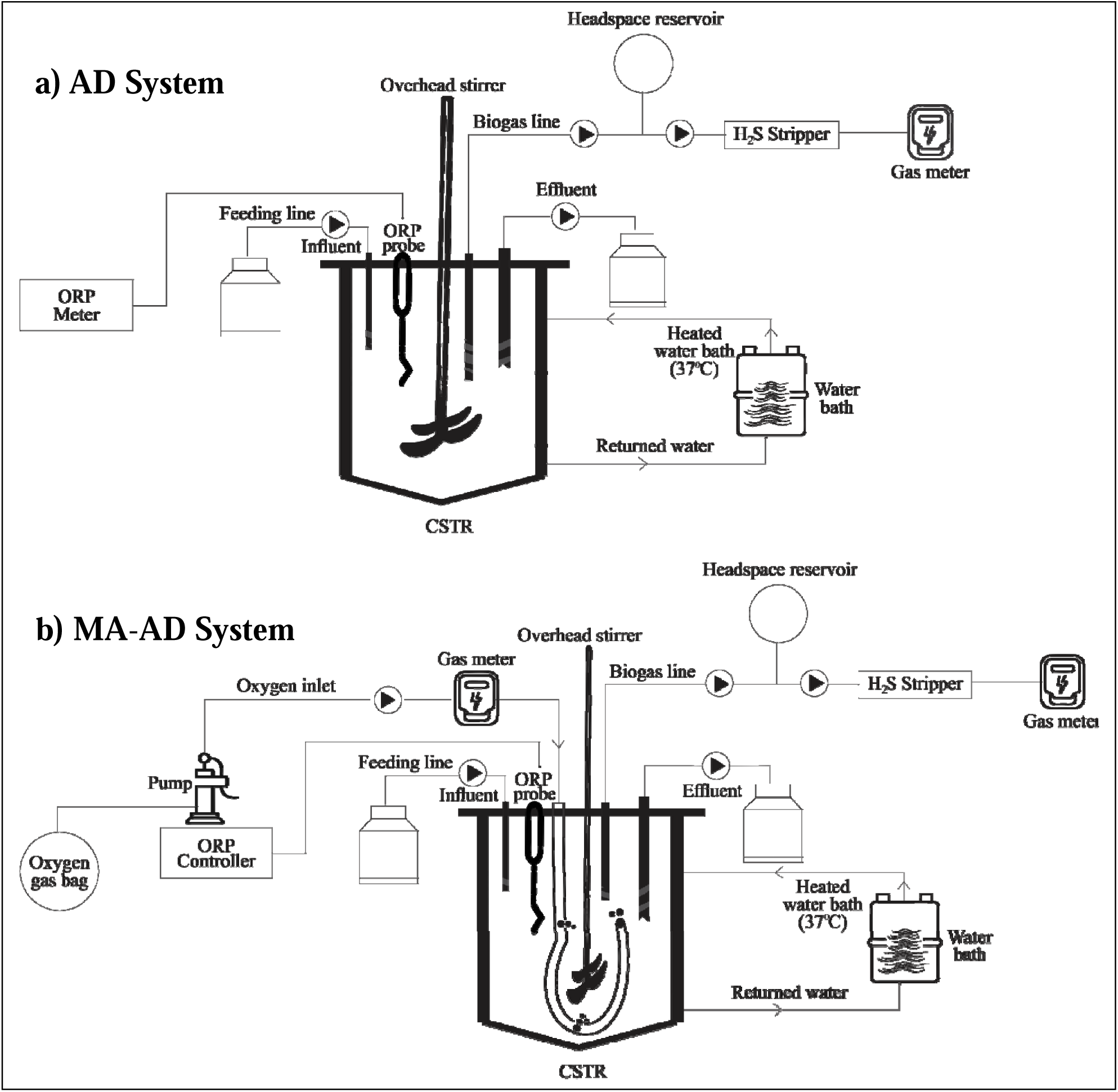
a) Anaerobic digestion (AD) reactor with oxidation-reduction potential (ORP) meter, and b) Micro-aerated anaerobic digestion (MA-AD) reactor with custom-designed oxidation-reduction potential (ORP) controller. CSTR: continuously-stirred tank reactor; H_2_S = hydrogen sulfide.

### 2.2 ORP-controlled micro-aeration system

In this study, the experimental reactor was controlled by an ORP controller to maintain micro-aerobic conditions by targeting ORP increments +20 mV higher than the baseline anaerobic ORP during each study phase (i.e., Phase 1 = +20 mV; Phase 2= +40 mV; Phase 3 = +60 mV). Micro-aeration was achieved by injecting oxygen at a controlled flow rate of 10 mL⋅min^-1^ using a peristaltic pump. The daily oxygen dosing volume was tracked using integrated flow meters, which measured the oxygen flow before it entered the reactor, ensuring precise and accurate daily oxygen measurements. The ORP probe was highly sensitive to DO, detecting even small amounts, with DO concentrations as low as 0.1 mg⋅L^-1^ being measurable. As oxygen was injected, the ORP increased, indicating a shift towards a more oxidative environment. The ORP-controlled micro-aeration system was fully automated, with oxygen injection cycles triggered when the ORP fell below the target value and ceasing once the target ORP was achieved. This system maintained stable micro-aerobic conditions in the reactor, with consistent fluctuations of ORP around the target set point. The ORP setpoint values and the average oxygen dosing amount during each experimental stage are presented in **Table 1**

**Table 1:** The operational parameters of micro-aeration during the experiment.

| Phase | Set ORP (mV) | Dosing techniques | Average oxygen dosing ($\text{mL O}_2 \cdot \text{d}^{-1} \cdot \text{L}_{\text{reactor}}^{-1}$) | Gas type | Duration (Days) |
| --- | --- | --- | --- | --- | --- |
| Zeroth | -490 | N/A | NA | N/A | 160 (Days 0-159) |
| First | -470 | Automatic, on-demand | $28.70 \pm 1.03$ | Air | 129 (Days 160-289) |
| Second | -450 | Automatic, on-demand | $100.70 \pm 6.90$ | Oxygen | 70 (Days 290-360) |
| Third | -430 | Automatic, on-demand | $228.60 \pm 3.92$ | Oxygen | 70 (Days 361-430) |

### 2.3 Inoculum and CSTR operation

Spent coffee grounds (SCG) were obtained from a dining hall at the University of Georgia, Athens, USA, steeped for 48 hours at room temperature, and filtered to remove solid particles before being used as a model CPW substrate. The two reactors were inoculated with anaerobic digestion sludge (ADS) sourced from a wastewater treatment plant (Gwinnett, GA USA). Reconstituted whole milk powder (RWM) was used in place of milk processing wastewater as a co-digestion substrate to provide additional organic content, buffering capacity, and nutrients. The final substrate was a mixture containing per liter: SCG, 700 mL; ADS, 285.6 mL; RWM, 6.9 g; yeast, 1 g; 7.2 mL of mineral stock; and 7.2 mL of trace element solution. The mineral stock contains the following (in g·L^-1^): FeCl□·4H□O, 370; MgCl□·6H□O, 120; KCl, 86.7; NH Cl, 26.6; and CaCl□·2H□O, 16.7. The trace element solution consists of (in g·L^-1^): COCl_2_·6H□O, 2; MnCl□·4H□O, 1.33; H_3_BO_3_ 0.38; ZnSO□·7H□O, 0.29; Na MoO□·2H□O, 0.17; and CuCl□·2H□O, 0.18. The reactors were maintained at a constant temperature of 37 ± 1°C and operated with a hydraulic retention time of 40 days at an OLR of 0.6 g COD⋅L^-1^⋅d^-1^. The influent composition comprised 20% SCG, 60% RWM, and 20% ADS (i.e., COD basis). The substrate compositions are shown in **Table 2**.

**Table 2:**
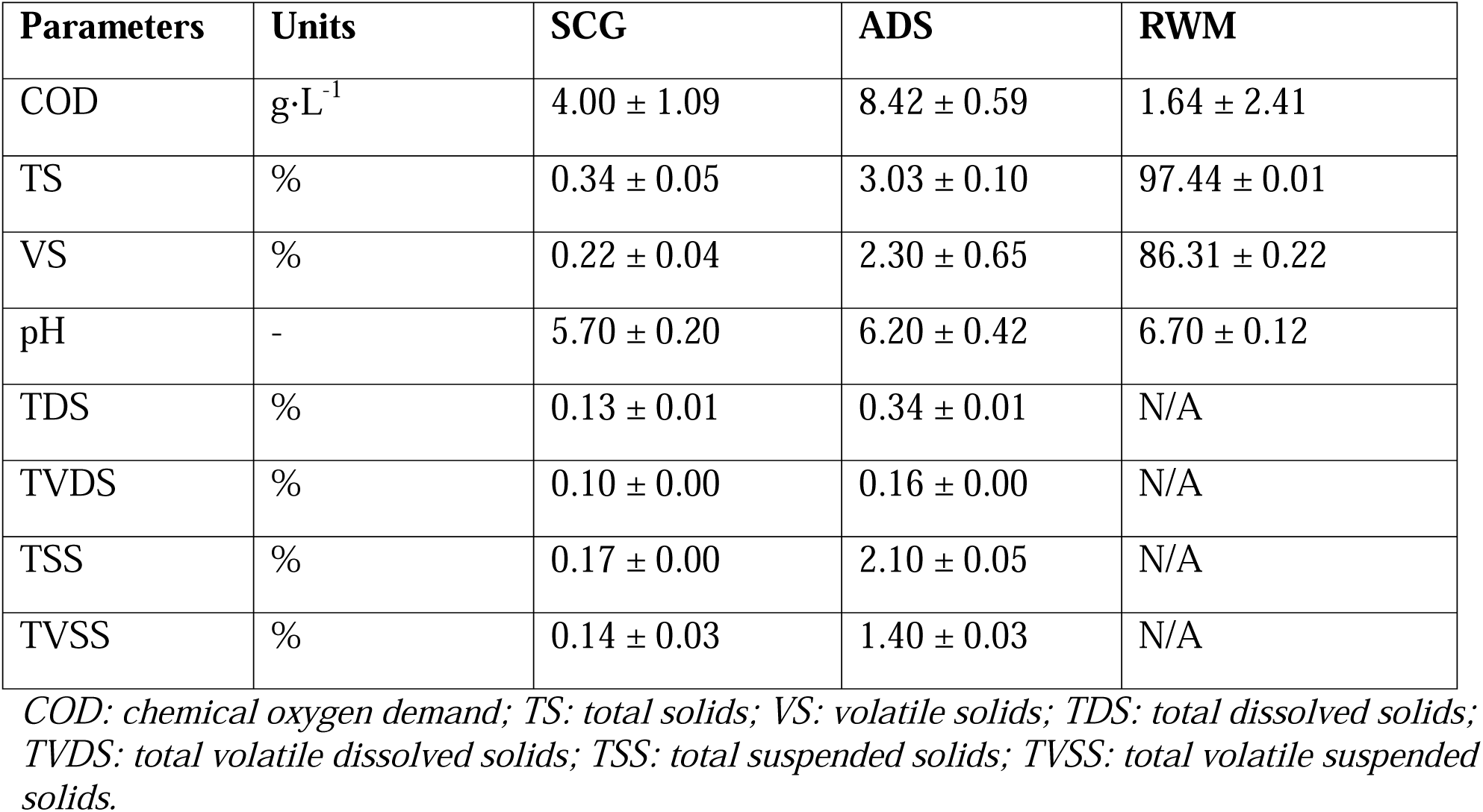
Substrate characterization data for spent coffee grounds (SCG), anaerobic digestion sludge (ADS), and reconstituted whole milk powder (RWM).

| Parameters | Units | SCG | ADS | RWM |
| --- | --- | --- | --- | --- |
| COD | $\text{g}\cdot\text{L}^{-1}$ | $4.00 \pm 1.09$ | $8.42 \pm 0.59$ | $1.64 \pm 2.41$ |
| TS | % | $0.34 \pm 0.05$ | $3.03 \pm 0.10$ | $97.44 \pm 0.01$ |
| VS | % | $0.22 \pm 0.04$ | $2.30 \pm 0.65$ | $86.31 \pm 0.22$ |
| pH | - | $5.70 \pm 0.20$ | $6.20 \pm 0.42$ | $6.70 \pm 0.12$ |
| TDS | % | $0.13 \pm 0.01$ | $0.34 \pm 0.01$ | N/A |
| TVDS | % | $0.10 \pm 0.00$ | $0.16 \pm 0.00$ | N/A |
| TSS | % | $0.17 \pm 0.00$ | $2.10 \pm 0.05$ | N/A |
| TVSS | % | $0.14 \pm 0.03$ | $1.40 \pm 0.03$ | N/A |

*COD: chemical oxygen demand; TS: total solids; VS: volatile solids; TDS: total dissolved solids; TVDS: total volatile dissolved solids; TSS: total suspended solids; TVSS: total volatile suspended solids.*

### 2.4 Analyses

Samples collected from the reactors were analyzed for pH, COD, total, dissolved, suspended, and volatile-solids (VS) following standard methods (APHA., 2012). Headspace gas samples were taken weekly to assess biogas composition (i.e., CH_4_, CO_2_), utilizing a gas chromatograph equipped with a methanizer and flame ionization detector (8610 C, SRI Instruments, Torrance, CA, USA). Nitrogen served as the carrier gas, with an inlet and detector temperature maintained at 110°C and a constant oven temperature set at 40°C. The concentrations of individual VFAs were determined using gas chromatography with a flame ionization detector (Agilent 6890, Santa Clara, CA, USA) according to the procedure described by Usack et al. (2014) with minor modifications. Hydrogen was used as the carrier gas, with the inlet temperature set to 200°C and the detector temperature set to 275°C. Individual VFA species were separated using a fused silica capillary column (NUKOL, 15 m × 0.53 mm × 0.50 μm film thickness; Supelco Inc., Bellefonte, PA, USA). The initial temperature was maintained at 70°C for 2 min, followed by an increase of 12°C/min until reaching 200°C, at which point it was held for an additional 2 min.

Turbidity was measured using a turbidity meter (HI 98703, Hanna Instruments, Woonsocket, RI, USA), following the manufacturer’s standard operating procedure. Chromophoric compounds were quantified using UV-Vis spectrophotometry (Cary 60, Agilent Technologies, Santa Clara, CA, USA) equipped with a diode array detector. Raw CPW was scanned from 200 to 400 nm, revealing a maximum absorbance at 270 nm. A calibration curve was established from serial dilutions of CPW, and effluent samples were analyzed accordingly. Caffeine was extracted using a liquid-liquid extraction method (Vandeponseele et al., 2021). The supernatants from centrifuged samples underwent three chloroform/methanol extractions, followed by solvent removal using a Büchi R-210 Rotavapor connected to a V-850 vacuum controller and a V-700 vacuum pump. The residue was resolubilized in HPLC-grade water, filtered (0.45-µm nylon), and analyzed via HPLC (Model 1100, Agilent Technologies, Santa Clara, CA, USA) with a variable wavelength detector. Separation was achieved using a reversed-phase Luna C18(2) column (4.6 × 250 mm, 5 µm particle size; Phenomenex, Torrance, CA, USA). An isocratic mobile phase was employed comprising water/methanol (65/35, v/v) with 1% acetic acid, a 1 mL·min^-^¹ flow rate, and a 20 µL injection volume. Caffeine was detected at 280 nm and quantified via calibration curves.

### 2.5 Statistical analysis

Statistical differences between the AD and MA-AD reactors were evaluated using one-way analysis of variance (ANOVA), followed by Tukey’s Honest Significant Difference (Tukey’s HSD) post-hoc test for pairwise comparisons. All analyses were conducted in R software (version 4.3.1). The stats package was used for ANOVA, and the agricolae and multicomp packages were employed for post-hoc tests and multiple comparisons. Data sets for methane yield, total solids (TS), VS, COD, and pH were assessed for normality and homogeneity of variance using the Shapiro-Wilk and Levene’s tests, respectively. Where assumptions were not met, appropriate transformations or non-parametric equivalents were applied. All statistical tests were performed at a significance level of α = 0.05. Data are presented as mean ± standard deviation unless otherwise stated.

## 3.0 Results and discussion

### 3.1 Reactor stabilization and microbial adaptation in micro-aerated anaerobic digestion

3.1.1 *pH dynamics and reactor stability*

The pH levels in the AD and MA-AD reactors were monitored throughout the study to assess the impact of micro-aeration on reactor stability. During the start-up phase (Phase 0), both reactors stabilized at a baseline pH of 7.08 ± 0.02 **(Figure 2),** indicating that the AD process was functioning normally and consistently under both conditions. During the stabilization phase, the main performance parameters (i.e., pH, biogas production, VFA concentration) varied less than +/- 10% of the running average, indicating consistent pseudo-steady-state operation between reactors (Usack et al., 2012). Initially, air was introduced as the dosing gas to facilitate micro-aeration in the MA-AD reactor. However, between Day 160 and Day 290, a reduction in CO partial pressure in the reactor headspace due to headspace dilution led to a gradual increase in pH. This pH shift was primarily caused by chemical changes related to gas-liquid CO equilibria rather than biological processes. Notably, this rise in pH did not correspond to any significant alterations in VFA accumulation or methane production. Pure oxygen gas was introduced on Day 280 to address the pH imbalance, replacing air. Following this intervention, a decrease in pH was observed in the MA-AD reactor compared to the AD reactor. This drop below baseline levels can be attributed to enhanced microbial activity under microaerobic conditions, promoting the production of organic acids by facultative microorganisms, as reported by Girotto et al. (2018). The switch from air to pure oxygen likely created more favorable conditions for the proliferation of acidogenic bacteria (Zhang et al., 2005), thereby intensifying acid production and maintaining higher levels of dissolved CO_2_.

**Figure 2:**
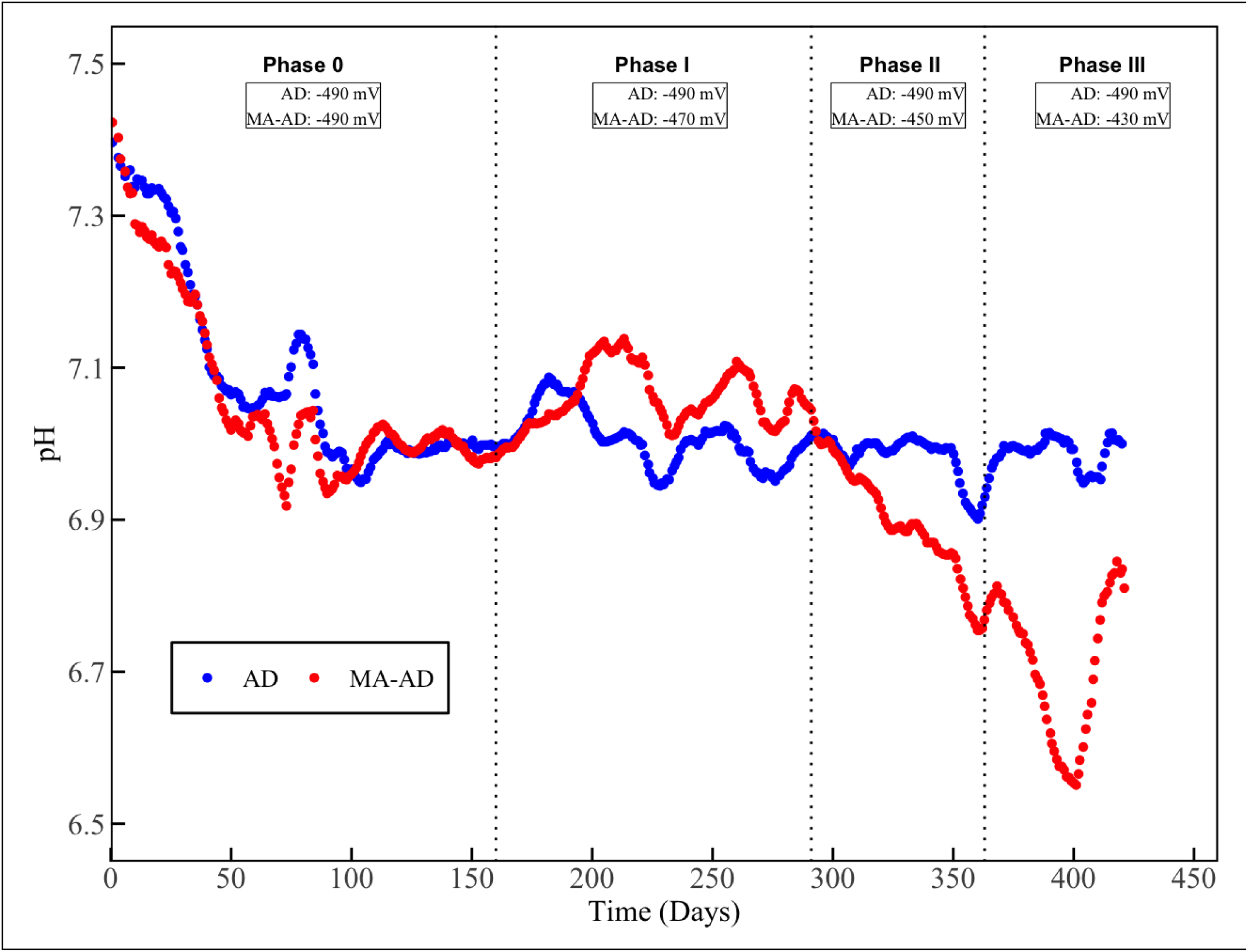
pH variation over time in the anaerobic digestion (AD) reactor and micro-aerated anaerobic digestion (MA-AD) reactor.

Throughout the study, caffeine was periodically introduced into the reactors to elevate its concentration, which, in turn, influenced the pH dynamics. On Day 230 (Phase I), both reactors were spiked with caffeine at a concentration of 200 mg·L□^1^ to achieve higher substrate levels and assess the effects of micro-aeration on caffeine degradation, as the steady-state caffeine concentrations in the reactor broths were low (i.e., 1.3 mg·L□^1^). This spiking event led to a marked drop in pH between Day 230 and Day 249, which could be attributable to 1) the inhibition of methanogenesis or 2) the microbial breakdown of caffeine into acidic metabolites such as theobromine, theophylline, paraxanthine, and various low-molecular-weight organic acids. The transient accumulation of these degradation products increased system acidity, resulting in a pH decline. A second caffeine spike of 200 mg·L□^1^ was administered between Day 352 and Day 369 (Phase II) to further assess the influence of micro-aeration on caffeine degradation. Consistent with the first spike, this intervention induced another drop in pH. Subsequently, between Day 403 and Day 411, a third caffeine spiking event led to an additional pH decrease in both reactors. Despite these drops, the reactors stabilized after a few days, particularly in the MA-AD reactor. From Day 395 onwards, the MA-AD reactor re-stabilized at 6.8, following an initial drop to 6.56 **(Figure 2)**.

#### 3.1.2 COD removal efficiency

The COD removal efficiencies were evaluated to assess the degradation of organic matter under anaerobic and micro-aerated conditions. Throughout the experimental phases, the total COD (TCOD) removal efficiency was 61.32 ± 17.98% for the AD reactor and 63.90 ± 17.29% for the MA-AD reactor **(Figure 3).** Similarly, soluble COD (SCOD) removal efficiencies were 88.29 ± 11.29% and 91.04 ± 7.04% for the AD and MA-AD reactors, respectively. Statistical analysis revealed no significant differences between the two reactors in TCOD removal (p = 0.58) and SCOD removal (p = 0.24). Furthermore, when COD removal efficiencies were analyzed for the experimental phases alone, the differences remained statistically insignificant (p = 0.413 for TCOD and p = 0.719 for SCOD). These findings suggest that the shift in acid production pathways, if any, due to micro-aeration neither enhanced nor impaired the overall degradation of organic matter. The results align with the findings of Duarte et al. (2024), who similarly reported no significant differences in SCOD removal between micro-aerated and non-micro-aerated reactors.

**Figure 3:**
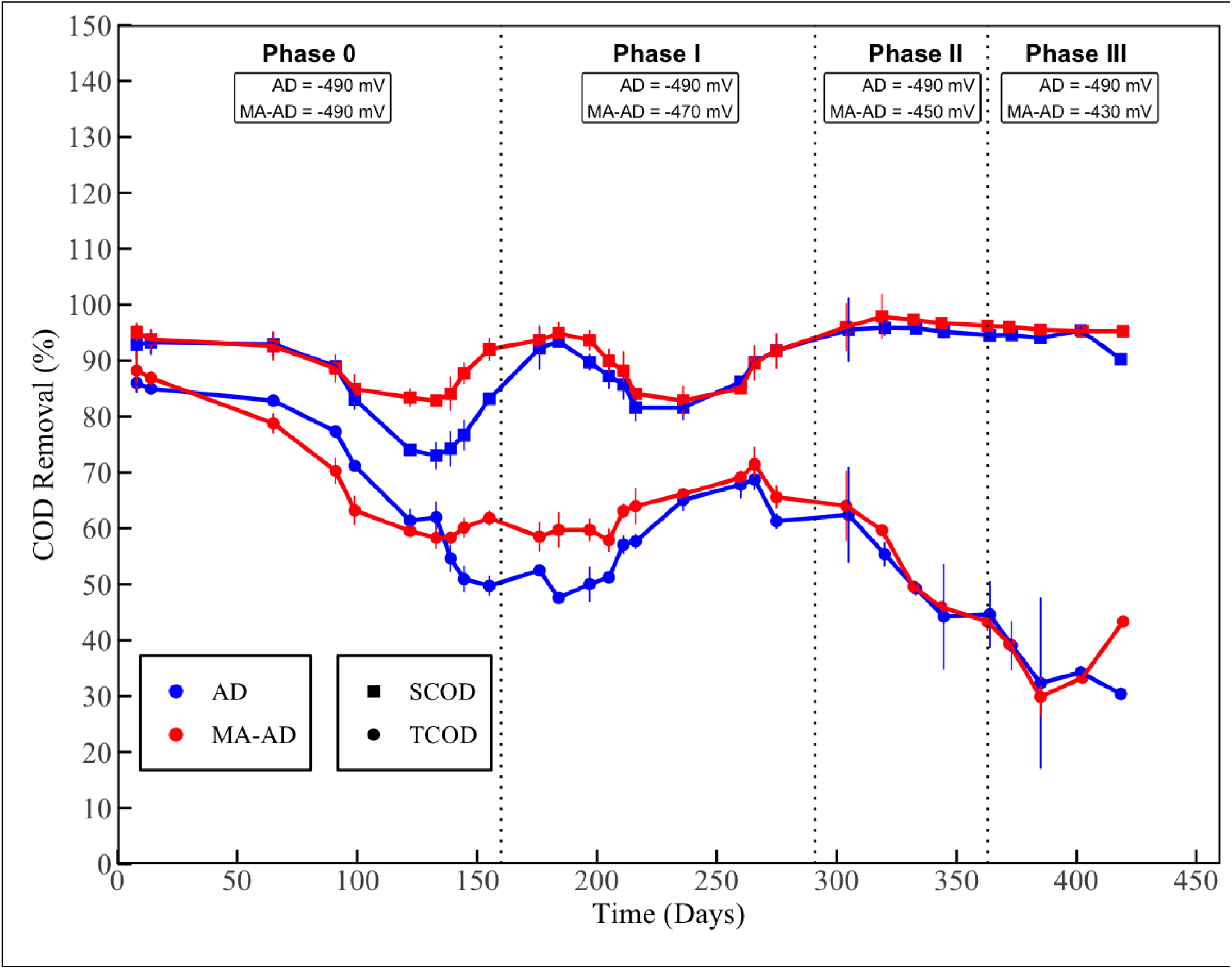
Chemical oxygen demand (COD) removal efficiency in the anaerobic digestion (AD) reactor and the micro-aerated anaerobic digestion (MA-AD) reactor. The vertical lines at each measurement point represent the standard deviation of the technical replicates. SCOD = soluble chemical oxygen demand; TCOD = total chemical oxygen demand.

An increase in hydrolysis rates typically causes an increase in the steady-state SCOD concentration. However, the MA-AD reactor exhibited a 2.85% lower average SCOD concentration than the AD reactor. This modest SCOD reduction combined with lower methane yields suggests microaeration had 1) no significant effect on hydrolysis rates or 2) increased hydrolysis rates, but the hydrolysis products were quickly metabolized (Harirchi et al., 2022), where they were assimilated into microbial biomass (Fernández-Domínguez et al., 2023) or fully oxidized to non-methanogenic end products (i.e., CO_2_, H_2_O) (Gaballah et al., 2025; Harirchi et al., 2022). Hence, the absence of elevated SCOD does not preclude an increase in hydrolytic activity under micro-aerobic conditions but could imply more efficient substrate turnover or a diversion of intermediates away from methane production.

#### 3.1.3 VFA concentrations and microbial activity

The VFA concentrations were closely monitored to evaluate the potential impacts of micro-aeration on acidogenesis and methanogenesis. VFA levels remained stable throughout the study, with concentrations of 364.80 ± 49.40 mg·L□^1^ in the MA-AD reactor and 383.90 ± 88.20 mg·L□^1^ in the AD reactor. Despite notable pH shifts caused by changes in gas dosing strategies, no significant VFA accumulation was observed in either reactor. Under micro-aerobic conditions, the system likely favored the production of non-VFA organic acids or other acidic intermediates, such as lactic acid, succinic acid, or similar compounds, which were not directly measured in this study. If micro-aeration stimulated VFA generation through increased acidogenic activity, it must have simultaneously increased VFA catabolism, as no VFA accumulation occurred. These findings are consistent with those reported by Canul Bacab et al. (2020), who observed comparable shifts in microbial metabolic pathways under oxygen-limited environments. Additionally, the low OLR used in this study likely contributed to maintaining reactor stability by limiting acetate availability for methanogenesis, which prevented VFA accumulation. This observation supports the findings of Fu et al. (2023), who reported that the influence of micro-aeration on VFA accumulation is closely linked to oxygen dosing rates and OLR.

### 3.2 Impact of micro-aeration on TS, VS, dissolved and suspended solids dynamics

#### 3.2.1 TS and VS reduction

In this study, TS and VS removal were evaluated under AD and MA-AD conditions. A significant difference in TS removal efficiency was observed between the two reactors, with the MA-AD reactor showing a 6.54 ± 7.11% lower reduction relative to the AD reactor **(Figure 4)**. On average, TS removal in the AD reactor was 51.78 ± 2.98%, whereas in the MA-AD reactor, it was 48.37 ± 4.36%. VS removal was also statistically significantly different, with the AD reactor achieving 63.08 ± 3.25% and the MA-AD reactor achieving 60.46 ± 4.00%, corresponding to an average 4.15 ± 3.86% lesser reduction in the MA-AD system relative to the AD system **(Figure 4)**. To gain more insight into these dynamics, each operational phase was analyzed separately using independent t-tests. TS removal showed a statistically significant difference in Phase I (p = 0.0103), while Phases II (p = 0.0976) and III (p = 0.0521) were not statistically significant, though the latter approached significance. For VS removal, the differences were not significant in Phase I (p = 0.4019) and Phase II (p = 0.3849) but became statistically significant in Phase III (p = 0.0090). This pattern suggests that while the initial impact of micro-aeration on VS degradation was limited, its effects became more apparent in later stages, potentially due to microbial adaptation or accumulation of biomass and partially degraded organics.

**Figure 4:**
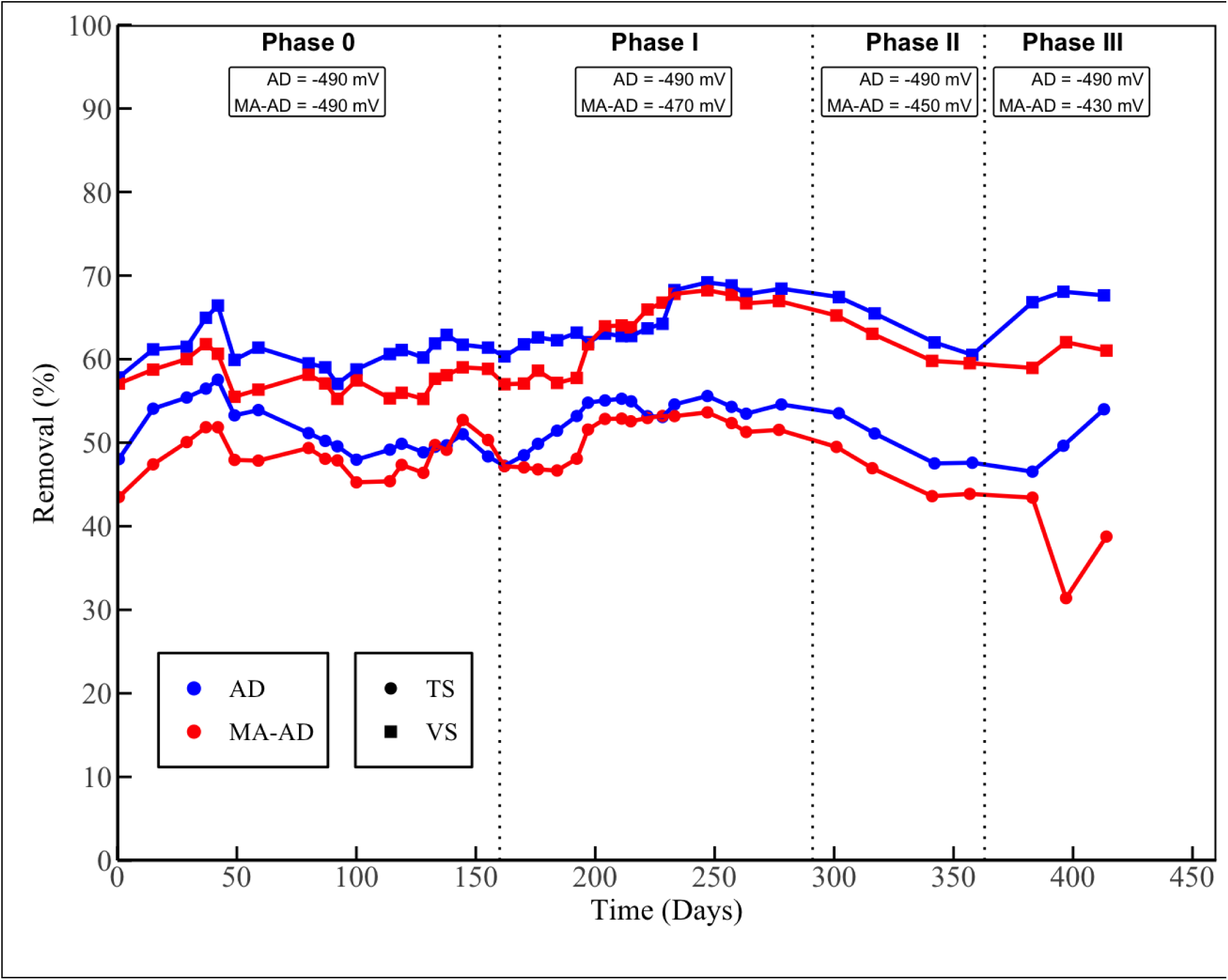
Total and volatile solids removal in the anaerobic digestion (AD) reactor and the micro-aerated anaerobic digestion (MA-AD) reactor.

The reduction in TS removal efficiency in the MA-AD reactor may be attributed to several factors, including an increase in biomass content resulting from heightened cellular growth stimulated by micro-aerobic conditions (Diak et al., 2013) and altered microbial dynamics. These shifts in microbial communities under limited oxygen exposure may have favored biomass accumulation over effective degradation of particulate matter, thereby reducing overall solids removal efficiency. Previous studies have reported that micro-aeration can alter the microbial community structure by enhancing the growth of facultative microorganisms that thrive under low-oxygen conditions (Morais et al., 2024), which could result in different metabolic pathways for the degradation of solids. These pathways, while potentially enhancing the production of organic acids or other metabolites, may not be as efficient at breaking down the bulk solids, including the more recalcitrant fractions of the solids. Specifically, Romero et al. (2021) reported that micro-aeration did not enhance TS removal from AD treatment of sewage sludge from municipal water resource recovery facilities.

Furthermore, this suggests that AD was as effective as MA-AD in breaking down the more biodegradable volatile components in the organic matter. Perhaps the advantages of micro-aeration are only realized when high concentrations of less degradable VS are present and not the more easily degradable VS. These results are consistent with findings from Diak et al. (2013), who reported that micro-aeration did not enhance the removal of TS and VS of primary sludge.

#### 3.2.2 Suspended solids and dissolved solids removal

Effluent samples were collected from the AD and MA-AD reactors at the end of each treatment phase and analyzed in triplicate to evaluate total suspended solids (TSS), total volatile suspended solids (TVSS), total dissolved solids (TDS), and total volatile dissolved solids (TVDS) concentrations. The TSS concentrations in the MA-AD reactor effluent were consistently higher than those observed in the AD reactor across all phases (**Figure 5 and Table 3**). Specifically, in Phase I, the MA-AD reactor exhibited a TSS concentration of 5.10 ± 0.20 g·L□^1^, while the AD reactor had 4.60 ± 0.10 g·L□^1^. In Phase II, the TSS concentrations were 5.80 ± 0.10 g·L□^1^ for MA-AD and 4.70 ± 0.30 g·L□^1^ for AD. In Phase III, a marked increase in TSS concentrations was observed in the AD and MA-AD reactors. Specifically, the AD reactor was 6.20 ± 0.46 g·L□^1^, and the MA-AD reactor was 7.6 ± 0.70 g·L□^1^. These differences were statistically significant (p < 0.05). This consistent upward trend in both treatment conditions suggests that micro-aeration was not the sole factor driving the TSS increase. While the higher final value in the MA-AD reactor may still point to an amplifying effect due to finite oxygen availability, the concurrent rise in the strictly anaerobic control indicates that other factors were also at play. These may include cumulative biomass accumulation over time, the progressive breakdown of bound or colloidal solids, or reduced sludge settleability due to prolonged reactor operation.

**Figure 5:**
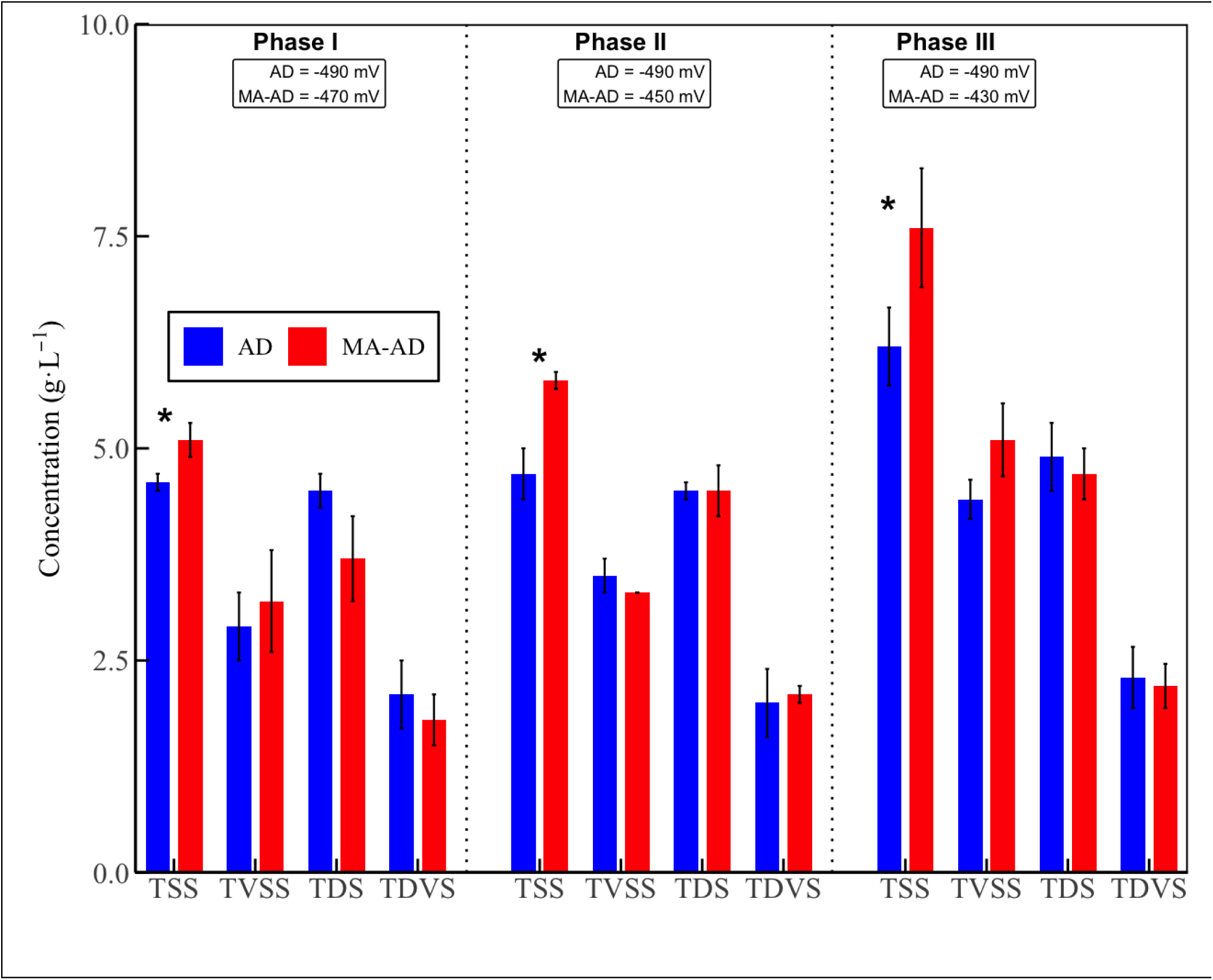
Total suspended solids (TSS), total volatile suspended solids (TVSS), total dissolved solids (TDS), and total dissolved volatile solids (TDVS) levels across treatment phases in the anaerobic digestion (AD) reactor and the micro-aerated anaerobic digestion (MA-AD) reactor.

**Table 3:**
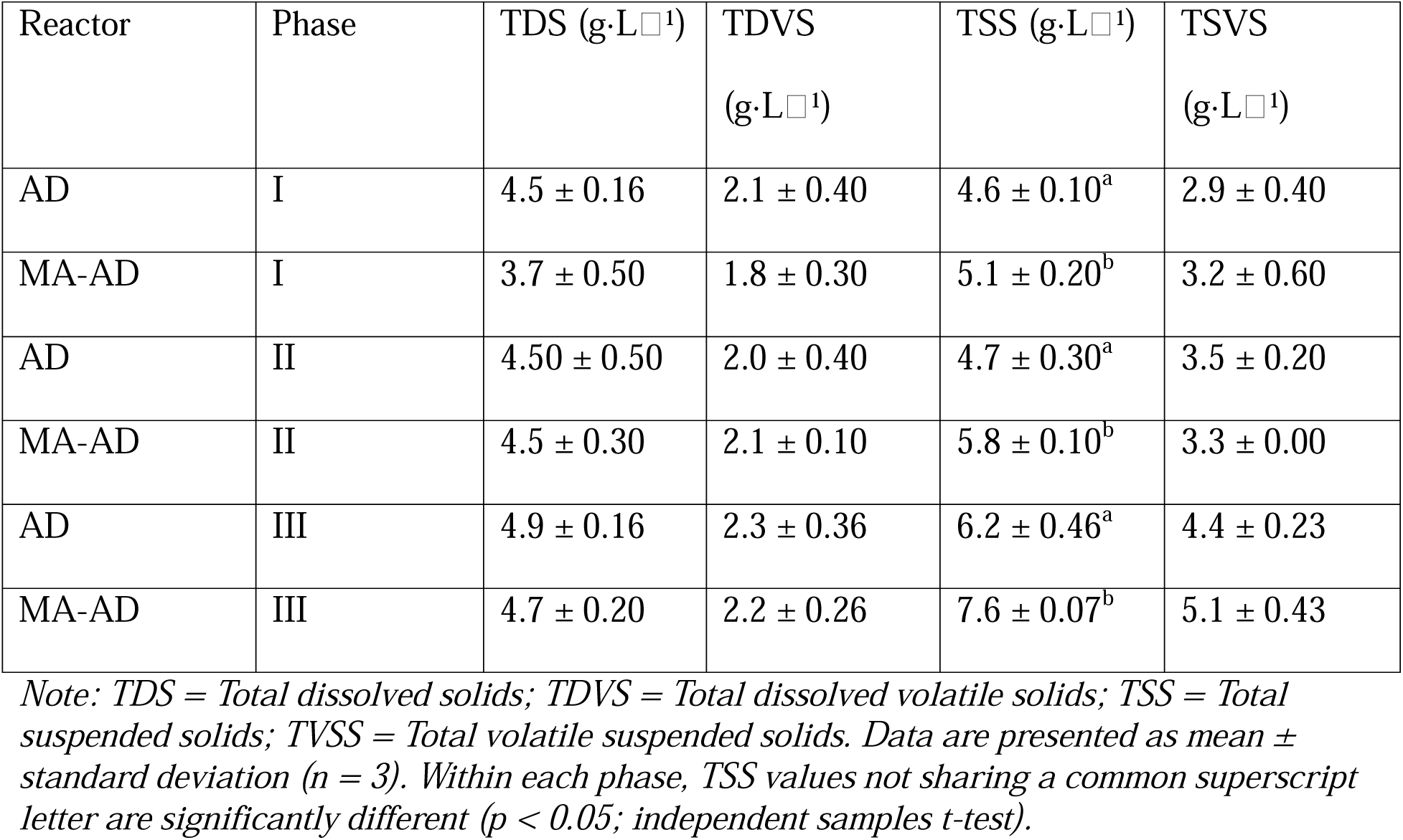
Effects of micro-aeration on dissolved and suspended solid concentrations across different phases.

| Reactor | Phase | TDS (g·L<sup>-1</sup>) | TDVS<br>(g·L<sup>-1</sup>) | TSS (g·L<sup>-1</sup>) | TSVS<br>(g·L<sup>-1</sup>) |
| --- | --- | --- | --- | --- | --- |
| AD | I | 4.5 ± 0.16 | 2.1 ± 0.40 | 4.6 ± 0.10<sup>a</sup> | 2.9 ± 0.40 |
| MA-AD | I | 3.7 ± 0.50 | 1.8 ± 0.30 | 5.1 ± 0.20<sup>b</sup> | 3.2 ± 0.60 |
| AD | II | 4.50 ± 0.50 | 2.0 ± 0.40 | 4.7 ± 0.30<sup>a</sup> | 3.5 ± 0.20 |
| MA-AD | II | 4.5 ± 0.30 | 2.1 ± 0.10 | 5.8 ± 0.10<sup>b</sup> | 3.3 ± 0.00 |
| AD | III | 4.9 ± 0.16 | 2.3 ± 0.36 | 6.2 ± 0.46<sup>a</sup> | 4.4 ± 0.23 |
| MA-AD | III | 4.7 ± 0.20 | 2.2 ± 0.26 | 7.6 ± 0.07<sup>b</sup> | 5.1 ± 0.43 |

*Note: TDS = Total dissolved solids; TDVS = Total dissolved volatile solids; TSS = Total suspended solids; TVSS = Total volatile suspended solids. Data are presented as mean ± standard deviation (n = 3). Within each phase, TSS values not sharing a common superscript letter are significantly different (p < 0.05; independent samples t-test).*

The elevated TSS levels in the MA-AD reactor further suggest complex interactions between suspended solids and microbial communities under micro-aerobic conditions **(Figure 5)**. The limited oxygen introduced likely promoted the growth of facultative anaerobes, which thrive in low-oxygen environments. Microbial biomass contributes to TVSS, and a numerical increase in TVSS was observed in Phase I and III within the MA-AD reactor relative to the AD reactor, suggesting an increase in biomass growth. This microbial shift may have influenced the flocculation and aggregation of solids, enhancing their retention within the reactor (Liu et al., 2023).

In addition to these biological factors, physicochemical mechanisms may have contributed to the observed increase in TSS under micro-aerobic conditions. Efendi et al. (2023) reported that increased aeration time and airflow in wastewater treatment systems resulted in a progressive increase in TSS, strongly correlated with DO levels. The study attributed this trend to the reaction between DO and dissolved metal ions, particularly Fe²□, forming insoluble Fe(OH) precipitates that increase measured TSS. In this study, Fe² was introduced as part of the trace mineral supplement in the feed. Although dosed at relatively low concentrations, the oxidative environment created by micro-aeration likely facilitated the conversion of Fe² to Fe³□, followed by precipitation as Fe(OH)□, thereby contributing directly to the elevated TSS values observed in the MA-AD reactor. This provides a plausible mechanistic explanation for the significant difference in TSS between the MA-AD and AD reactors in both treatment phases.

However, Zouari and Al Jabiri (2015) reported contrasting results, noting that micro-aeration improved the digestibility of TSS by enhancing the breakdown of complex organic matter, thus preventing its accumulation in sludge. However, such outcomes may be highly system-dependent, varying with substrate composition, oxygen exposure regimes, and microbial consortia. Furthermore, Jenicek et al. (2011) observed no significant differences in TSS or TVSS concentrations during the micro-aerobic treatment of digested sludge, further suggesting that micro-aeration may not universally enhance solids degradation. Similarly, Uman et al. (2018) observed no significant difference in solids concentrations when applying the Fenton reaction (i.e., H_2_O_2_ [oxidizer] + Fe^2+^) to pretreat wastewater biosolids before AD.

In contrast, TDS, TVDS, and TVSS removal efficiencies showed no statistically significant differences between the AD and MA-AD reactors across treatment phases (**Figure 5**, **Table 3**). These closely aligned values across systems and phases suggest that micro-aeration did not significantly enhance the microbial breakdown of either dissolved or particulate solids. This finding supports previous reports that strictly anaerobic conditions may be more effective for degrading VS due to the dominance of specialized obligate anaerobes that metabolize VFA and other organics (Appels et al., 2008). In summary, while micro-aeration influenced the behavior of suspended solids, as is evident from the elevated TSS in the MA-AD reactor, it did not provide a measurable advantage in removing either dissolved or suspended VS.

#### 3.2.3 Turbidity reduction and effluent clarity improvement

The impact of micro-aeration on turbidity was evaluated by monitoring changes in Nephelometric Turbidity Units (NTU) across two treatment phases. Turbidity serves as an indirect indicator of suspended colloidal particles and potential microbial or physicochemical destabilization within the reactor. Under strictly AD conditions, turbidity values were recorded at 2030 ± 77 NTU in Phase I and 2404 ± 162 NTU in Phase II **(Figure 6)**. In contrast, under MA-AD conditions, turbidity increased to 2143 ± 89 NTU in Phase I and significantly more in Phase II, reaching 3147 ± 114 NTU. In Phase III, this trend became even more pronounced, with turbidity rising to 2526 ± 138 NTU in the AD reactor and sharply to 6064 ± 242 NTU in the MA-AD reactor **(Figure 6)**. These findings suggest the redox conditions and the aeration rate influence turbidity.

**Figure 6:**
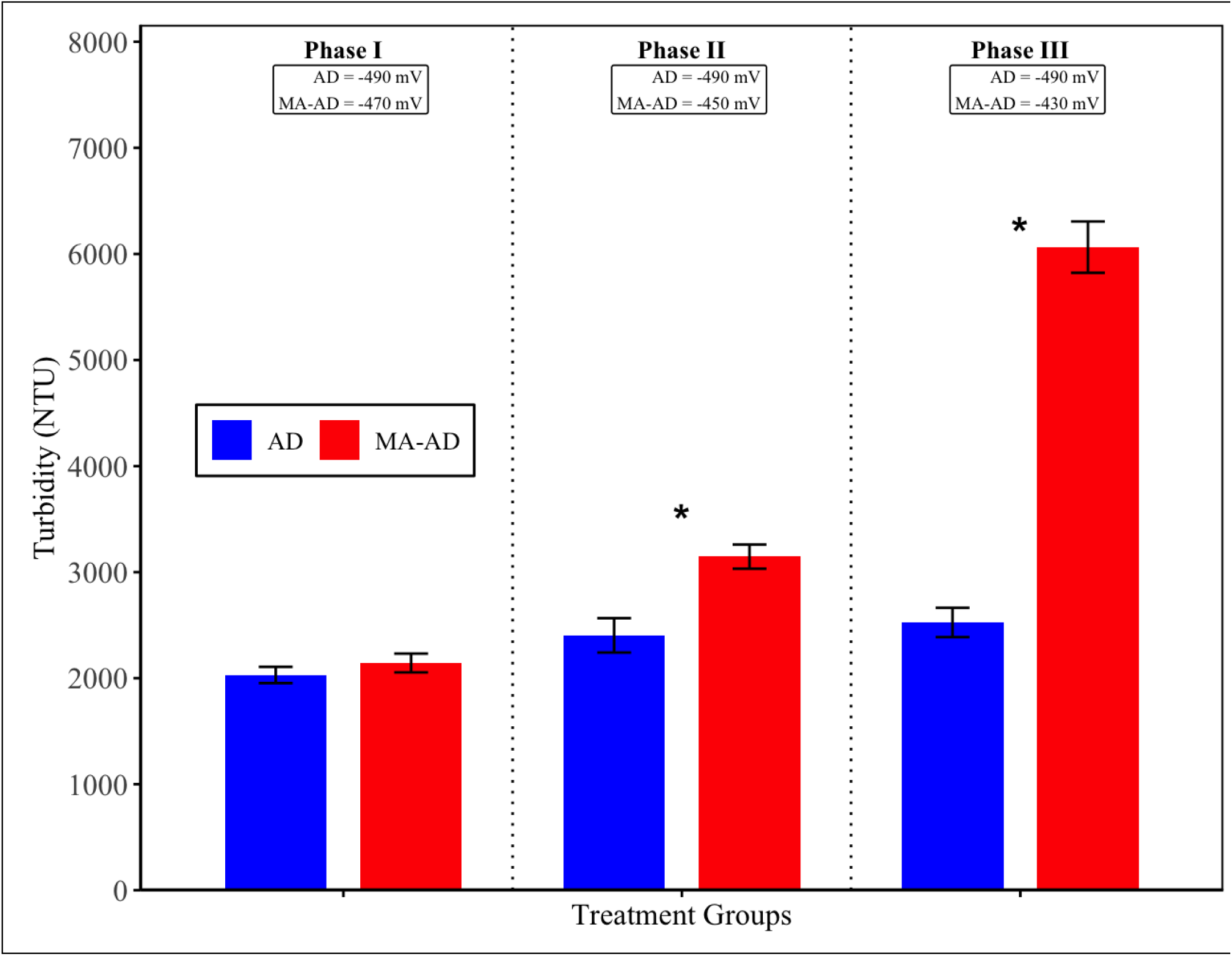
Turbidity levels across the treatment phases in the anaerobic digestion (AD) reactor and the micro-aerated anaerobic digestion (MA-AD) reactor.

The higher redox potential and oxygen availability could alter microbial metabolism and destabilize floc structures by disrupting floc integrity, releasing fine particulate matter into suspension, thereby increasing turbidity (Zhang et al., 2019). It is also plausible that elevated microbial metabolic activity partially degraded solids into smaller, light-scattering particles, contributing to increased turbidity. Furthermore, under micro-aerobic conditions, physicochemical processes such as oxidation of dissolved sulfur compounds or metals could have contributed to turbidity. For example, Tang et al. (2004) showed that aeration in hydrogen sulfide-rich systems can lead to the formation of insoluble sulfur precipitates, which significantly increase turbidity. The high turbidity recorded in the MA-AD reactor during Phase III (6000 ± 240 NTU) suggests a threshold beyond which excessive oxygen input or prolonged aeration may induce floc disintegration and increase the release of fine particles. This supports the hypothesis that micro-aeration may trigger similar precipitation pathways in AD, particularly in the presence of sulfur- or metal-containing ions. In wastewater with high organic loads and complex compounds such as CPW, the effect of micro-aeration on turbidity removal may be limited unless specific factors, such as aeration position and oxygen content, are optimized (Zhang et al., 2019). In conclusion, while micro-aeration can positively influence microbial and physicochemical dynamics within AD systems, it also poses a risk of increasing turbidity, particularly under high oxygen dosing conditions. These results highlight the importance of fine-tuning the aeration strategy to balance biological enhancement with the potential destabilization of solids.

### 3.3 Methane yield in micro-aerobic digestion

Methane production was consistently higher in the AD reactor across all experimental phases of the study. During the start-up phase (Phase 0), the AD reactor produced approximately 100.0 ± 16.7 mL CH_4_⋅gCOD^-1^⋅L^-1^⋅d^-1^, while the MA-AD reactor yielded a comparable 101.9 ± 21.5 mL CH_4_⋅gCOD^-1^⋅L^-1^⋅d^-1^ **(Figure 7)**, indicating near-equal performance. Notably, the low methane yield observed during the start-up phase in the AD and MA-AD reactors shows the inherently low biodegradability of the co-digestion substrate: waste-activated sludge (WAS). Several studies have reported low methane yields from untreated WAS, typically ranging from 80 to 192 mL CH_4_⋅gVS^-1^, primarily due to its high concentration of degradation-resistant microbial cells, the low content of non-cellular biodegradable organic matter, and the general need for pretreatment to enhance hydrolysis and microbial accessibility (Feki et al., 2020; Guo et al., 2023; Kampioti & Komilis, 2022; Uman et al., 2018; Wang et al., 2016).

**Figure 7:**
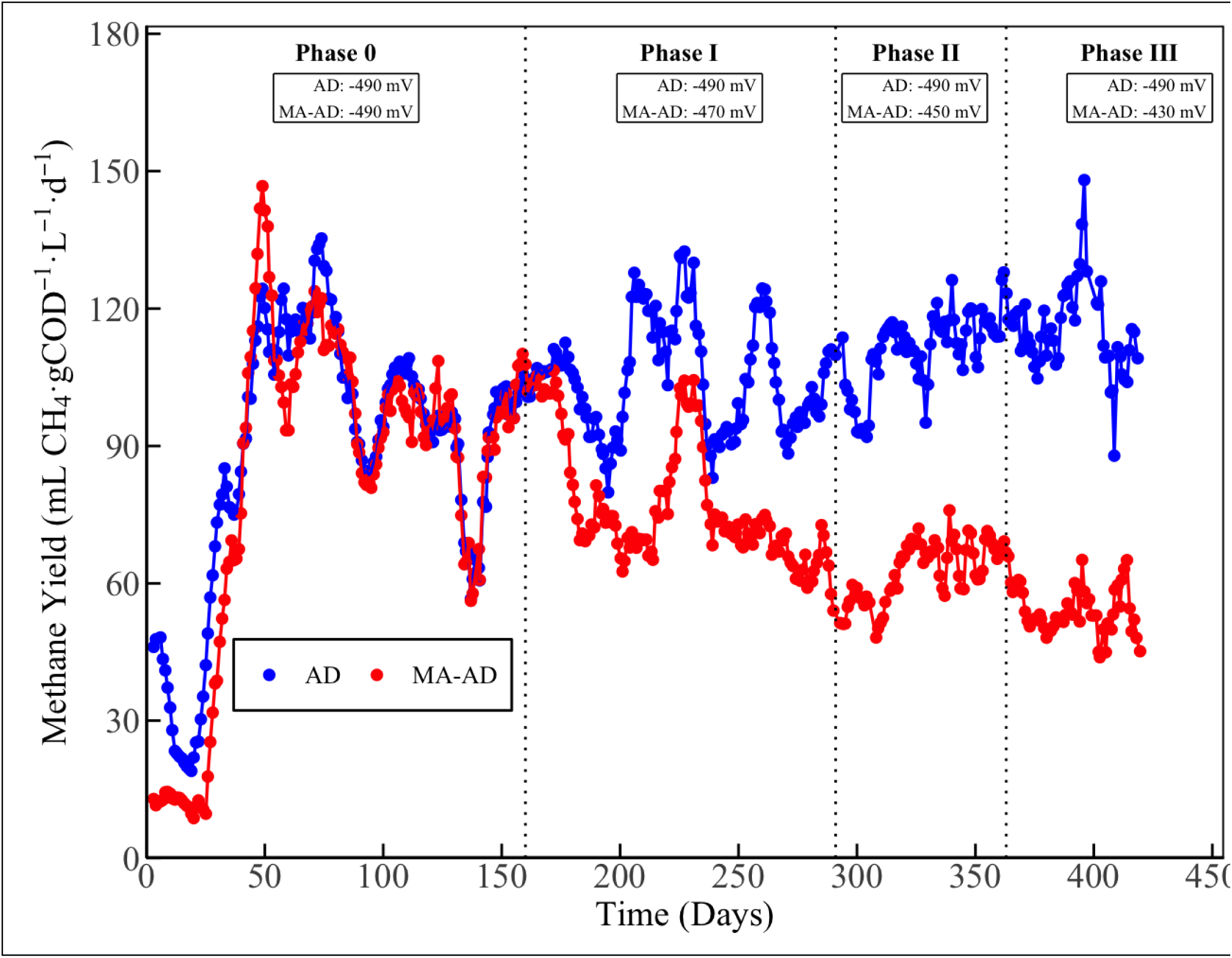
Methane yield in the anaerobic digestion (AD) reactor and micro-aerated anaerobic digestion (MA-AD) reactor.

This shows that co-digestion of WAS with CPW, another recalcitrant substrate, did not substantially enhance methane yield, especially in the absence of thermal, chemical, or enzymatic pretreatment. Additionally, Widjaja et al. (2017) reported that co-digesting coffee pulp without appropriate pretreatment does not significantly enhance methane yields due to the persistence of lignocellulosic and polyphenolic compounds. These results imply that co-digestion may not be a sufficient strategy to promote recalcitrant compound degradation in CPW; pretreatments may be essential to improve biodegradability and enhance methane recovery. In contrast, studies involving AD of readily degradable substrates such as food waste or its co-digestion with energy-rich materials like paper waste, cattle manure, or water hyacinth have consistently reported much higher methane yields, typically ranging from 388 to 607 mL CH_4_⋅gVS^-1^ (Kim & Oh, 2011; Marañón et al., 2012; Oduor et al., 2022; Usack & Angenent, 2015; Zhang et al., 2013), showing enhanced biodegradability and the limitations of using recalcitrant substrates such as WAS and CPW without pretreatment.

During the experimental phases, the methane production in the MA-AD reactor dropped to 77.5 ± 13.7 mL CH_4_⋅gCOD^-1^⋅L^-1^⋅d^-1^ in Phase I, a 23.9% reduction relative to 102.3 ± 12.1 mL CH_4_⋅gCOD^-1^⋅L^-1^⋅d^-1^ in the AD reactor. This decline became more pronounced in Phase II, where methane production in the MA-AD reactor decreased further to 62.8 ± 6.9 mL CH_4_⋅gCOD^-1^⋅L^-1^⋅d^-1^, representing a 43.5% decrease relative to the AD reactor, which generated 111.2 ± 8.3 mL CH_4_⋅gCOD^-1^⋅L^-1^⋅d^-1^. In Phase III, the reduced methane production continued, with the AD reactor producing 110.7 ± 19.47 mL CH□⋅gCOD□^1^ L□^1^ d^-1^, while methane production in the MA-AD reactor declined further to just 53.4 ± 6.03 mL CH□⋅gCOD□^1^□L□^1^ d□^1^ a reduction of nearly 52% compared to the control. The progressive decrease in methane yield under microaerobic conditions can be attributed to the oxygen sensitivity of methanogenic archaea. These organisms are obligate anaerobes, and their enzymatic systems are irreversibly inhibited by even trace amounts of oxygen (Botheju & Bakke, 2011; Girotto et al., 2018). The presence of oxygen alters redox balance and suppresses methanogenic pathways that are critical for converting VFA and hydrogen into methane. In addition to direct oxygen toxicity, several studies have shown that introducing oxygen into anaerobic reactors can shift the microbial community in favor of facultative and aerobic bacteria (Nguyen & Khanal, 2018). While such organisms may enhance the hydrolysis and acidogenesis stages of digestion, they compete directly with methanogens for substrates, particularly VFAs. If not properly managed, substrate competition can significantly reduce methane production (Nguyen, 2018; Yoda et al., 1987). As Nguyen et al. (2019) reported, effective micro-aeration demands careful regulation of oxygen loading to maintain the delicate balance between facultative bacteria and obligate anaerobes.

In the context of this study, the reduction in methane production may also be linked to the chemical complexity of the feedstock. Although CPW comprised 20% of the total COD in the co-digestion mix, the presence of recalcitrant compounds such as polyphenols, tannins, and caffeine may have contributed to localized or synergistic inhibitory effects on methanogenesis, particularly when combined with oxygen exposure in the MA-AD reactor. Methanogens are especially vulnerable to polyphenolic compounds, and even low concentrations can significantly impair their activity (Puchalska et al., 2021). Teng et al. (2024) investigated the effects of tea-derived polyphenols on methane production in cattle and found that even low doses suppressed methane output by altering the diversity and structure of the enteric microbial communities. This provides mechanistic support for the hypothesis that the polyphenolic content of CPW could inhibit methanogenesis, even at relatively low inclusion levels in the feedstock. Supporting these findings, literature reports on the AD of coffee-related wastes reflect similar methane yields. For instance, Latif et al. (2022) reported a yield of 14.797 ± 0.01 mL CH□⋅g^-1^ VS^-1^ during the co-digestion of coffee waste with thickened sludge. In a related study, Semaan et al. (2023) reported a methane yield of 119.7 mL CH□⋅g^-1^ VS^-1^ for AD of SCG. Additionally, Teixeira et al. (2024) reported 188 mL CH□⋅g^-1^ VS^-1^ from industrial SCG co-digested with food waste at high concentrations (75%), where inhibitory effects were observed. At a lower concentration (25%), however, the methane yield increased significantly to 351 mL CH□⋅g^-1^ VS^-1^.

Operational factors also played a role in the observed results. Between Days 130 and 150, methane production was abnormally low in both reactors due to construction-related adjustments that required intermittent reactor shutdown and reconfiguration. These data were excluded from the final methane yield analysis to preserve the integrity of inter-phase comparisons. A further factor that may have contributed to the observed reduction in methane production is the progressive increase in oxygen demand within the MA-AD reactor. Initially, volumetric oxygen dosing averaged 28.73 ± 1.10 mL O ⋅d^-1^⋅L_reactor_^-1^ (Phase I), but over time, the required oxygen input rose to as high as 100.73 ± 6.90 mL O ⋅d^-1^⋅L_reactor_^-1^ (Phase II) and further to 228.60 ± 3.92 mL O ⋅d^-1^⋅L_reactor_^-1^ (Phase III) (excluding Day 404 onwards) to maintain target redox conditions **(Figure 8)**. This substantial increase in oxygen dosage could indicate either a shift in microbial community oxygen consumption or reduced oxygen transfer efficiency. In either case, the elevated oxygen levels may have exacerbated the inhibition of obligate methanogenic archaea, as sustained or excessive oxygen exposure is known to suppress methane-forming pathways and disrupt anaerobic syntrophy. Additionally, oxygen introduction can favor the proliferation of facultative and aerobic microorganisms, leading to increased competition for key intermediates such as VFA, which are otherwise consumed by methanogens. This competition further reduces the availability of substrates for methane production and may hinder the stability of the methanogenic stage. At Day 404 (**Figure 8),** a sharp decline in oxygen dosing was observed shortly after caffeine addition. This decline may have been triggered by the caffeine spike, which could have temporarily inhibited microbial respiration, particularly among facultative aerobes, thereby reducing oxygen uptake and prompting the ORP controller to lower oxygen supply. Alternatively, the observed pattern may reflect a combination of microbial adaptation, reduced overall metabolic oxygen demand, or natural stabilization of redox conditions near the ORP setpoint, requiring less oxygen input to maintain control.

**Figure 8:**
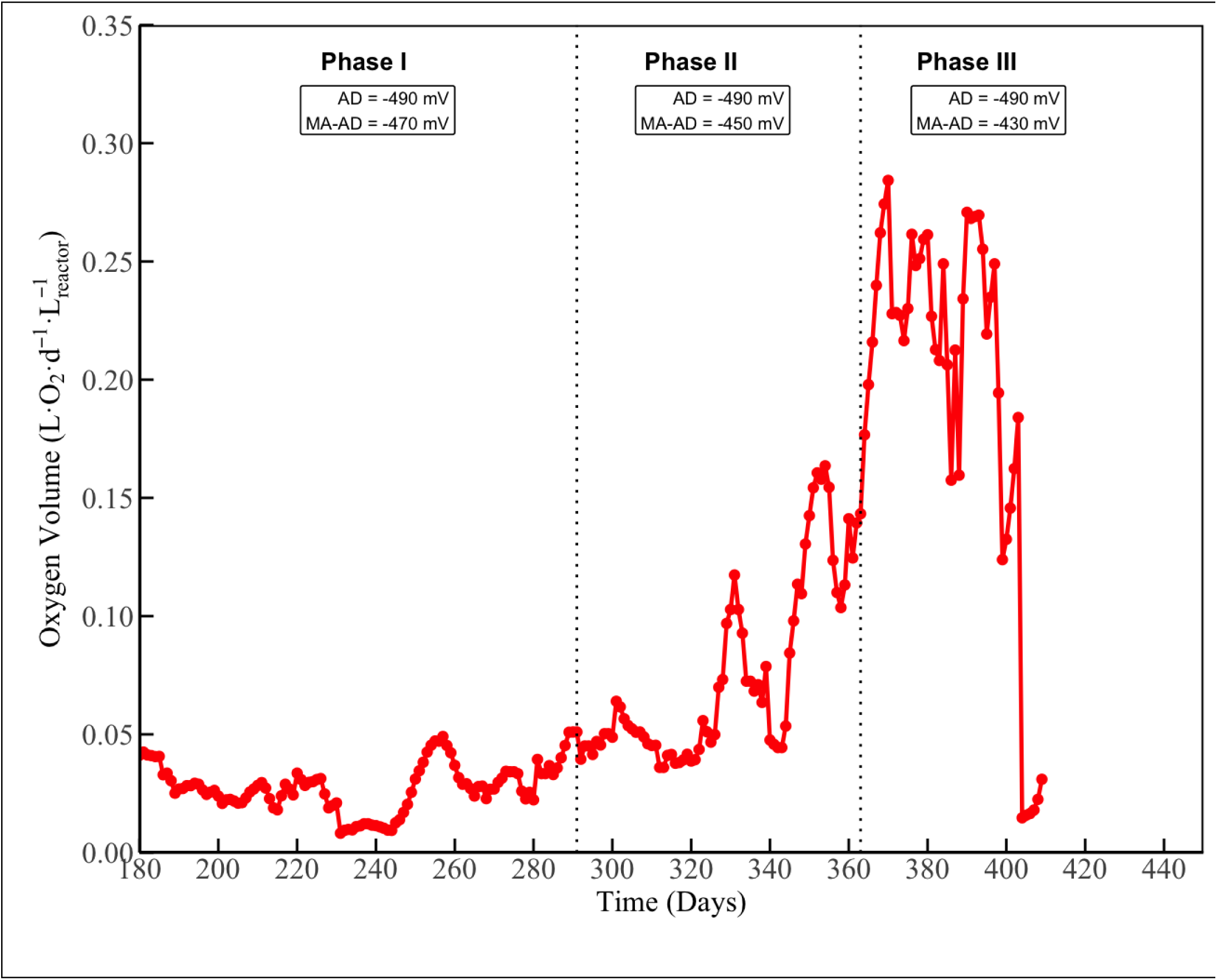
Oxygen dosing rate in the anaerobic digestion (AD) reactor and the micro-aerated anaerobic digestion (MA-AD) reactor.

Overall, while micro-aeration can stimulate microbial activity and accelerate the breakdown of complex organics, it may prove counterproductive when methane generation is the primary objective (Nguyen, 2018). Introducing oxygen can disrupt the anaerobic microbial ecosystem, inhibit methanogens, and dilute the biogas. These effects are particularly critical in systems where biogas yield and composition are a priority. In contrast, micro-aeration can be an effective and economical enhancement strategy when the operational focus shifts toward controlling intermediate products such as VFAs or enhancing hydrolysis (Nguyen, 2018). Ultimately, optimizing aeration intensity and controlling redox dynamics are essential for aligning system performance with specific treatment goals.

### 3.4. Degradation of recalcitrant and chromophoric compounds under micro-aerobic conditions

This study investigated the degradation behavior of recalcitrant compounds (caffeine and other chromophoric organics) under strictly AD and MA-AD conditions. This section provides insights into how oxygen exposure and redox dynamics influence the removal efficiency of aromatic and phenolic substances commonly present in CPW.

#### 3.4.1 UV spectral trends of chromophoric compounds under micro-aerobic conditions

Ultraviolet (UV) absorbance analysis was employed to quantify the relative concentration of chromophoric organic compounds in the effluents of AD and MA-AD reactors, using raw CPW as a calibration reference. An initial full-spectrum scan of the undiluted CPW revealed a maximum absorbance at 270 nm, within the wavelength range for polyphenols, aromatic rings, and other conjugated chromophores commonly present in agro-industrial waste streams. Serial dilutions of CPW were prepared to construct a calibration curve, from which the relative concentrations of UV-absorbing compounds in reactor effluents were back-calculated.

Effluent samples collected at the end of Phases I–III showed varied levels of residual chromophoric compounds. In Phase I, the relative concentration was 30.09 ± 1.13% in the AD reactor and 19.53 ± 0.82% in the MA-AD reactor, indicating slightly better removal under micro-aerobic conditions **(Figure 9).** However, in Phase II, the concentration in the MA-AD reactor spiked to 92.46 ± 3.06%, compared to 39.67 ± 1.94% in the AD reactor, suggesting a substantial decline in the degradation of chromophoric compounds under increased oxygen input. In Phase III, the MA-AD reactor again showed a higher residual concentration (70.11 ± 2.32%) compared to the AD reactor (42.51 ± 1.90%), showing that oxygen dosing at higher levels may impair the removal of UV-absorbing species. These trends are consistent with the findings from caffeine degradation trends discussed in Section 3.4.2, where the MA-AD reactor exhibited enhanced early-stage degradation kinetics but failed to achieve complete removal in later stages. This parallel behavior suggests a shared mechanistic limitation, potentially involving disruption of obligate anaerobic pathways or an oxygen-induced metabolic shift that favored partial oxidation over complete degradation.

**Figure 9:**
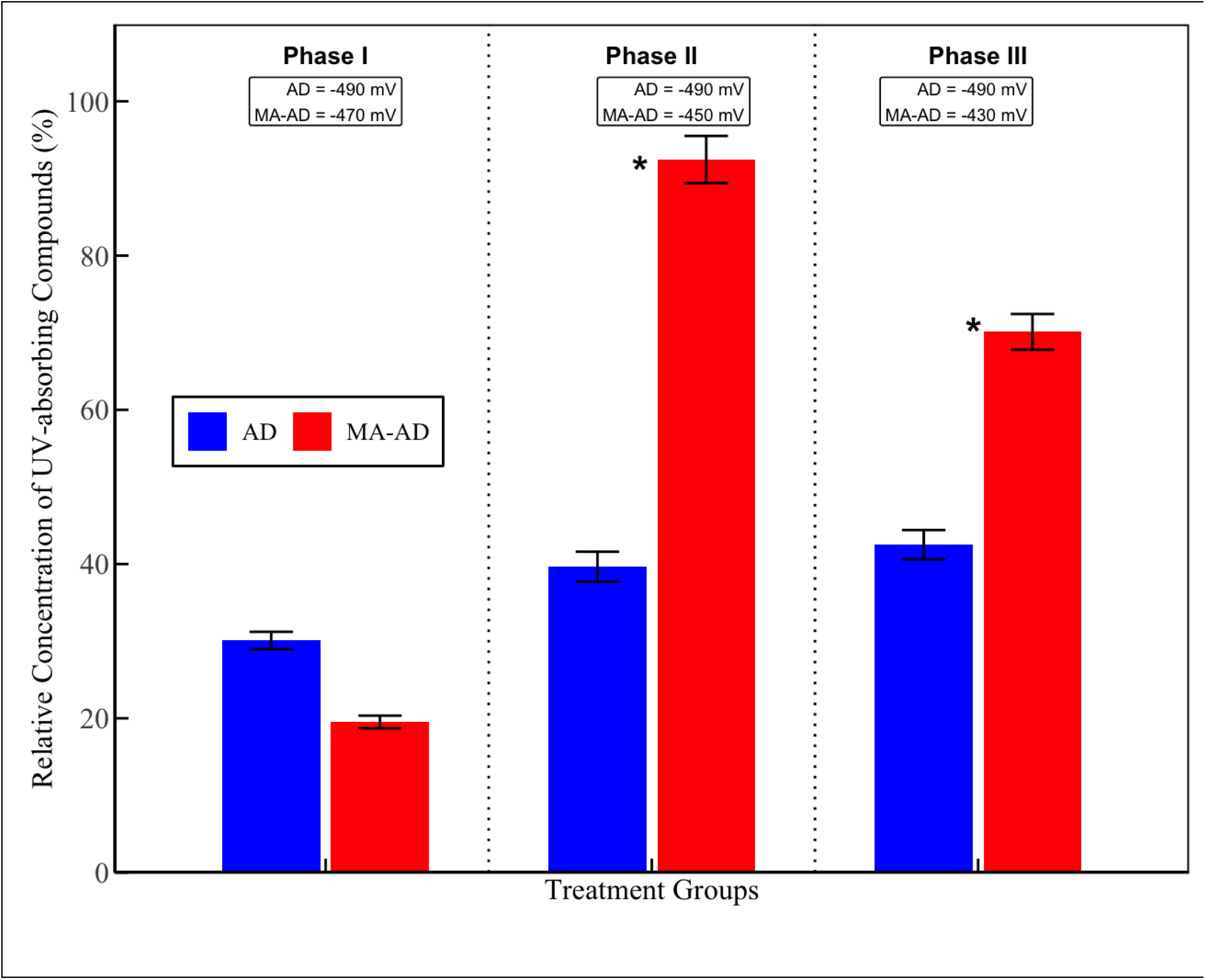
Removal of UV-absorbing compounds in the anaerobic digestion (AD) reactor and the micro-aerated anaerobic digestion (MA-AD) reactor.

This shows the critical importance of balanced oxygen input and microbial community dynamics in micro-aerated systems. While low-level micro-aeration may stimulate the oxidative breakdown of complex organics, excessive oxygen exposure appears to interfere with downstream anaerobic processes responsible for complete degradation. These findings emphasize the need for precise ORP control and tailored aeration strategies when applying micro-aeration to treat aromatic-rich waste streams like CPW.

#### 3.4.2 Caffeine Degradation Performance

Caffeine degradation was evaluated over three independent spiking stages using an initial concentration of ∼200 mg⋅L^-1^ of caffeine in the reactors. The reactors were operated under ORP-controlled conditions, with oxygen supplied as needed to the MA-AD reactor based on the set ORP. For Phase I, the MA-AD reactor showed faster caffeine degradation compared to the AD reactor. Within 28 hours, caffeine concentrations decreased by >85% in the MA-AD reactor, while the AD reactor achieved approximately 75% removal **(Figure 10).** This initial enhancement is likely due to the stimulation of facultative microbial activity, which has been shown to accelerate the breakdown of aromatic compounds under oxygen-limited conditions (Aydin et al., 2025). Facultative microorganisms capable of co-metabolizing caffeine through oxidative demethylation may have been enriched under these conditions, contributing to the observed advantage. However, by 36 hours, both reactors had reached near-complete degradation (∼90%), indicating convergence of long-term removal efficiency regardless of oxygen input. A similar trend was reported by Chen et al. (2018), who demonstrated that complete caffeine degradation can be achieved under strictly anaerobic conditions using an anaerobic membrane reactor, reporting a long-term removal efficiency of 87.5 ± 5.3%. In their study, caffeine degradation followed a two-step methanogenic pathway: the initial hydrolysis of caffeine into intermediates, followed by conversion of those intermediates into VFAs, and eventually CH_4_ and CO□. Notably, the transformation of hydrolysis products into VFAs was identified as the rate-limiting step of the process. Hydrolysis may have been rate limiting in the present study, given the lower SCOD concentration in the MA-AD reactor, and assuming a faster rate of hydrolysis product metabolism.

**Figure 10:**
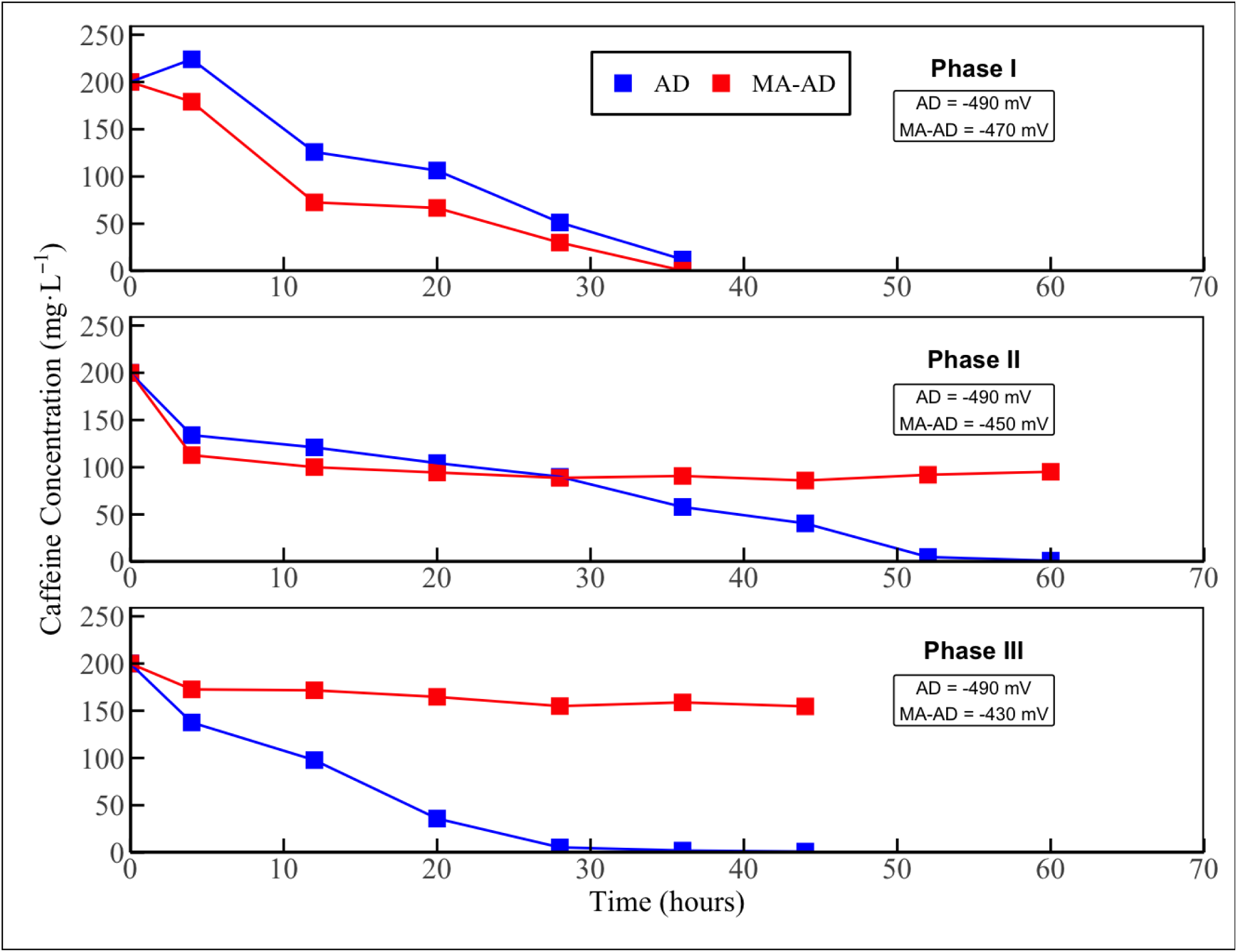
Caffeine degradation curve in the anaerobic digestion (AD) reactor and micro-aerated anaerobic digestion (MA-AD) reactor.

In Phase II, both reactors were spiked again after reaching baseline levels. Once more, the MA-AD reactor demonstrated a faster initial degradation rate than the strictly AD reactor. By 28 hours, the caffeine concentration in the MA-AD reactor had decreased to 94.4 ± 0.2 mg⋅L^-1^, compared to 104.4 ± 0.3 mg⋅L^-1^ in the AD reactor **(Figure 10).** This early-phase degradation advantage supports earlier observations that micro-aeration may enhance the initial transformation of caffeine, likely due to the stimulation of facultative microbial populations capable of oxidative co-metabolism. However, this initial performance advantage was not sustained. In Phase II, beyond 36 hours, the trend reversed. The AD reactor showed continued degradation, reaching near-complete removal (∼1.05 mg⋅L^-1^) by 84 hours, while caffeine levels in the MA-AD reactor stagnated between 85–95 mg⋅L^-1^ from 36 hours onward **(Figure 10)**. Furthermore, throughout this stage, the volume of oxygen dosed into the MA-AD reactor increased markedly compared to earlier phases **(Figure 8)**. This increase may reflect shifts in microbial oxygen demand or reduced microbial responsiveness to redox levels. Such excessive oxygenation could inhibit obligate anaerobes, impairing the conversion of caffeine intermediates into VFAs and methane (Lu & Imlay, 2021).

In Phase III, both reactors were again spiked with 200 mg⋅L^-1^ of caffeine. Caffeine removal in the MA-AD reactor remained incomplete, with concentrations only declining from 200 mg⋅L^-1^ to 172.68 ± 3.38 mg⋅L□^1^ after 4 hours and finally plateauing at 132.36 ± 0.08 mg⋅L□^1^ after 60 hours, like the trend observed in Phase II. Meanwhile, the AD reactor showed consistent degradation, reaching near-complete removal (0.95 ± 0.38 mg⋅L□^1^) by 44 hours. This may imply that the microbial population in the MA-AD reactor had shifted away from specific microbial species capable of degrading caffeine or that an accumulation of intermediate metabolites from previous stages interfered with degradation. Alternatively, increased oxygen pressure may have limited the activity of facultative anaerobes without fully re-establishing anaerobic syntrophy.

Although aerobic microorganisms generally exhibit greater metabolic efficiency and higher energy yields than their anaerobic counterparts, their ability to degrade caffeine in the MA-AD reactor may have been constrained by several factors. The microbial community present may not have been fully adapted to aerobic caffeine degradation, requiring specific enzymes or co-metabolic conditions that were not prevalent in the MA-AD reactor. Furthermore, oxygen toxicity may have inhibited certain microbial groups that play a crucial role in caffeine breakdown, disrupting enzymatic pathways and leading to metabolic inefficiencies. Additionally, aerobic pathways may lead to the accumulation of intermediate metabolites that were not readily degraded under the given conditions, potentially inhibiting further microbial activity. These findings indicate the importance of oxygen dosing precision and microbial community acclimation in maintaining the effectiveness of micro-aerated systems for recalcitrant compound degradation.

### 4.0 Conclusion and future perspective

This study comprehensively evaluates MA-AD in treating CPW, focusing on its effects on reactor stability, solids removal, turbidity, methane yield, and the degradation of inhibitory compounds such as caffeine. The results demonstrate that while micro-aeration can enhance early-stage microbial activity, hydrolysis, and degradation rates of certain recalcitrant compounds, its long-term success depends on carefully balanced operational parameters. Critically, oxygen dosing emerges as a paramount factor in the effective operation of MA-AD reactors. When applied in low, controlled quantities, oxygen can stimulate facultative microbial activity, promote partial oxidation of complex substrates, and accelerate the initial breakdown of polyphenolic compounds such as caffeine. However, excessive oxygen introduced unintentionally through overcompensated redox control can disrupt anaerobic microbial consortia, inhibit key methanogenic pathways, and diminish methane yields. This highlights the importance of finely tuned ORP regulation and real-time monitoring to prevent oxygen overload. Furthermore, the trace mineral composition of the substrate plays a pivotal role in reactor behavior. In this study, the presence of Fe² and other micronutrients may have catalyzed physicochemical reactions such as floc formation and particulate precipitation, influencing turbidity and solids retention. These interactions underscore the interconnectedness between substrate chemistry and reactor performance, especially under shifting redox levels.

Inhibitory effects from phenolic-rich substrates such as CPW were also evident, with results suggesting that even at relatively low concentrations (20% of total COD), compounds such as tannins and caffeine can exert significant microbial stress, particularly under oxidative stress. The observed shifts in pH and microbial efficiency further reinforce the need for cautious substrate formulation and reactor adaptation. Pretreatment strategies, such as alkaline hydrolysis, enzymatic conditioning, or strategic co-digestion, may offer viable approaches to enhance substrate bioavailability and mitigate inhibition. As demonstrated in comparable studies, these methods can substantially improve methane yields and solids reduction when dealing with lignocellulosic or polyphenol-laden waste streams. Ultimately, the success of MA-AD hinges on the integration of operational control (oxygen and ORP), feedstock chemistry (trace elements and inhibitors), and process design (OLR, mixing, and pretreatment). Future research should aim to define optimal oxygen loading thresholds, assess microbial community shifts with high-resolution sequencing, and deploy advanced analytical tools (e.g., LC-MS, metabolomics) to track degradation intermediates and byproducts. When appropriately managed, MA-AD holds significant promise for the efficient and stable treatment of complex industrial wastewater such as that generated in coffee processing.

## Abbreviations

AD BOD: Anaerobic digestion Biochemical oxygen demand
CPW: Coffee processing wastewater
COD: Chemical oxygen demand
DO: Dissolved oxygen
NTU: Nephelometric turbidity units
MA-AD: Micro-aerated anaerobic digestion
ORP: Oxidation-reduction potential
TS: Total solids
TSS: Total suspended solids
TVSS: Total volatile suspended solids
VFAs: Volatile fatty acids
VS: Volatile solids

## Declarations

### Ethics approval and consent to participate

Not applicable.

### Consent for publication

Not applicable.

### Availability of data and materials

All data generated or analyzed during this study are included in this published article.

### Author contributions

KJT: Visualization, Data Curation, Formal Analysis, Conceptualization, Investigation, Methodology, Validation, Writing - Original Draft, Writing - Review & Editing. SOO: Sample Analysis, Writing - Review & Editing. WLK: Writing - Review & Editing. RBP: Writing - Review & Editing. JHS: Writing - Review & Editing. JGU: Supervision, Conceptualization, Project Administration, Writing - Review & Editing, Funding Acquisition.

## Acknowledgment

The first author would like to acknowledge Elizabeth Ziabtchenko, Ibrahim Bello, Shreya Riswadkar, and Nithya Sree Kotha for their help running the reactors.

